# Lead compounds for the development of SARS-CoV-2 3CL protease inhibitors

**DOI:** 10.1101/2020.08.03.235291

**Authors:** Sho Iketani, Farhad Forouhar, Hengrui Liu, Seo Jung Hong, Fang-Yu Lin, Manoj S. Nair, Arie Zask, Yaoxing Huang, Li Xing, Brent R. Stockwell, Alejandro Chavez, David D. Ho

## Abstract

We report the identification of three structurally diverse compounds – compound 4, GC376, and MAC-5576 – as inhibitors of the SARS-CoV-2 3CL protease. Structures of each of these compounds in complex with the protease revealed strategies for further development, as well as general principles for designing SARS-CoV-2 3CL protease inhibitors. These compounds may therefore serve as leads for the basis of building effective SARS-CoV-2 3CL protease inhibitors.

## Main Text

As the etiologic agent of COVID-19, SARS-CoV-2 has resulted in hundreds of thousands of deaths and caused rampant economic damage worldwide^1,2^. While some treatments have been identified, their clinical efficacy is low, making continued research essential^3,4^. Similar to other coronaviruses, SARS-CoV-2 encodes an essential 3CL protease (M_pro_) that processes its polyproteins, which has garnered interest as a target for potential viral inhibitors^5,6^. Here, we describe a series of compounds with inhibitory activity against the SARS-CoV-2 3CL and determine their structures in complex with the protease. These data provide general insights into the design of 3CL protease inhibitors, along with potential avenues by which these classes of compounds can be further developed.

We hypothesized that previously identified SARS-CoV-1 3CL protease inhibitors may also be effective against the SARS-CoV-2 3CL, given the conservation between the two proteases (96% amino acid identity)^1,2^. Using a biochemical assay to report SARS-CoV-2 3CL protease activity (**Extended Data Fig. 1a, b**), we identified three diverse compounds of interest: compound 4^7^, GC376^8^, and MAC-5576^9^, which had IC_50_ values (mean ± s.d.) of 0.149 ± 0.002 µM, 0.139 ± 0.016 µM, and 0.088 ± 0.003 µM, respectively (**Fig. 1a, b**). Encouraged by these results, we then tested these compounds for inhibition of SARS-CoV-2 viral replication. We found that compound 4 and GC376 could block viral infection (EC_50_ values (mean ± s.d.): 3.023 ± 0.923 µM and 4.481 ± 0.529 µM, respectively), whereas MAC-5576 did not (**Fig. 1c**). Finally, we confirmed that these compounds did not result in cytotoxicity to the cells at the tested concentrations (**Extended Data Fig. 2**).

**Fig 1.**
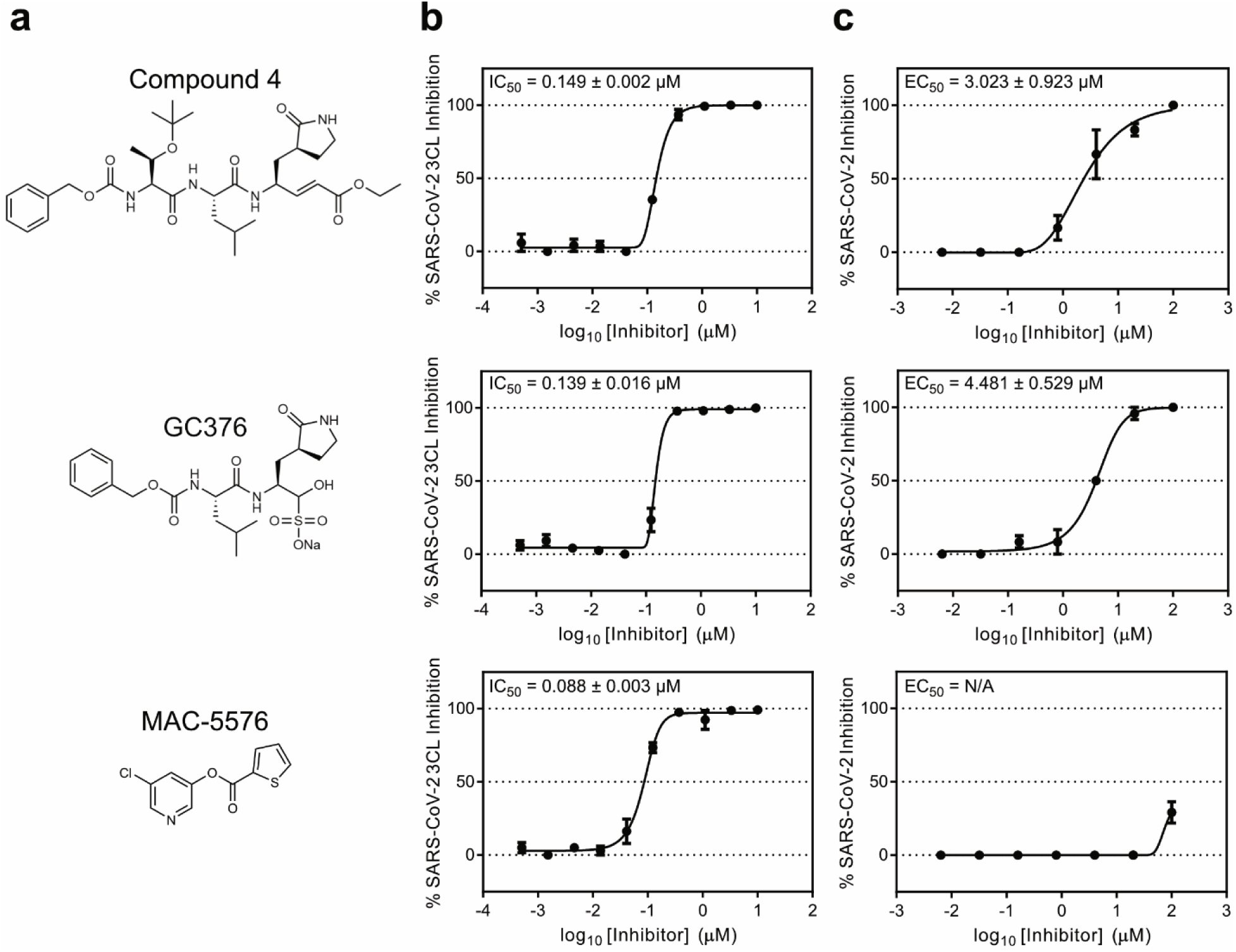
Inhibition of SARS-CoV-2 3CL protease by compound 4, GC376, and MAC-5576. **a**, Chemical structures of the three compounds in this study. **b**, Inhibition of purified native SARS-CoV-2 3CL protease by each compound. **c**, Inhibition of SARS-CoV-2 viral replication by each compound. Data are shown as mean ± s.e.m. for two or three technical replicates. IC_50_ and EC_50_ values are indicated as mean ± s.d. for two biological replicates.

As the three compounds exhibited inhibitory activity against the SARS-CoV-2 3CL, we proceeded to solve the crystal structure of the *apo* 3CL protease alone and of each of these compounds in complex with the protease to understand their mechanism of binding as well as to guide future structure-based optimization efforts. We note that while MAC-5576 did not exhibit activity in the cellular assay, its low molecular weight and reasonable biochemical activity prompted us to pursue its crystallization as well, as our goal was to broadly investigate inhibitory scaffolds for the SARS-CoV-2 3CL protease. Crystals were obtained (see **Methods** for detailed information) and structures at 1.85 Å, 1.80 Å, 1.83 Å, and 1.73 Å resolution limits for *apo* 3CL and 3CL bound to compound 4, GC376, and MAC-5576, respectively, were solved (**Fig. 2a, b, c, Extended Data Fig. 3, and Extended Data Fig. 4;** see **Supplementary Table 1** for statistics).

**Fig 2.**
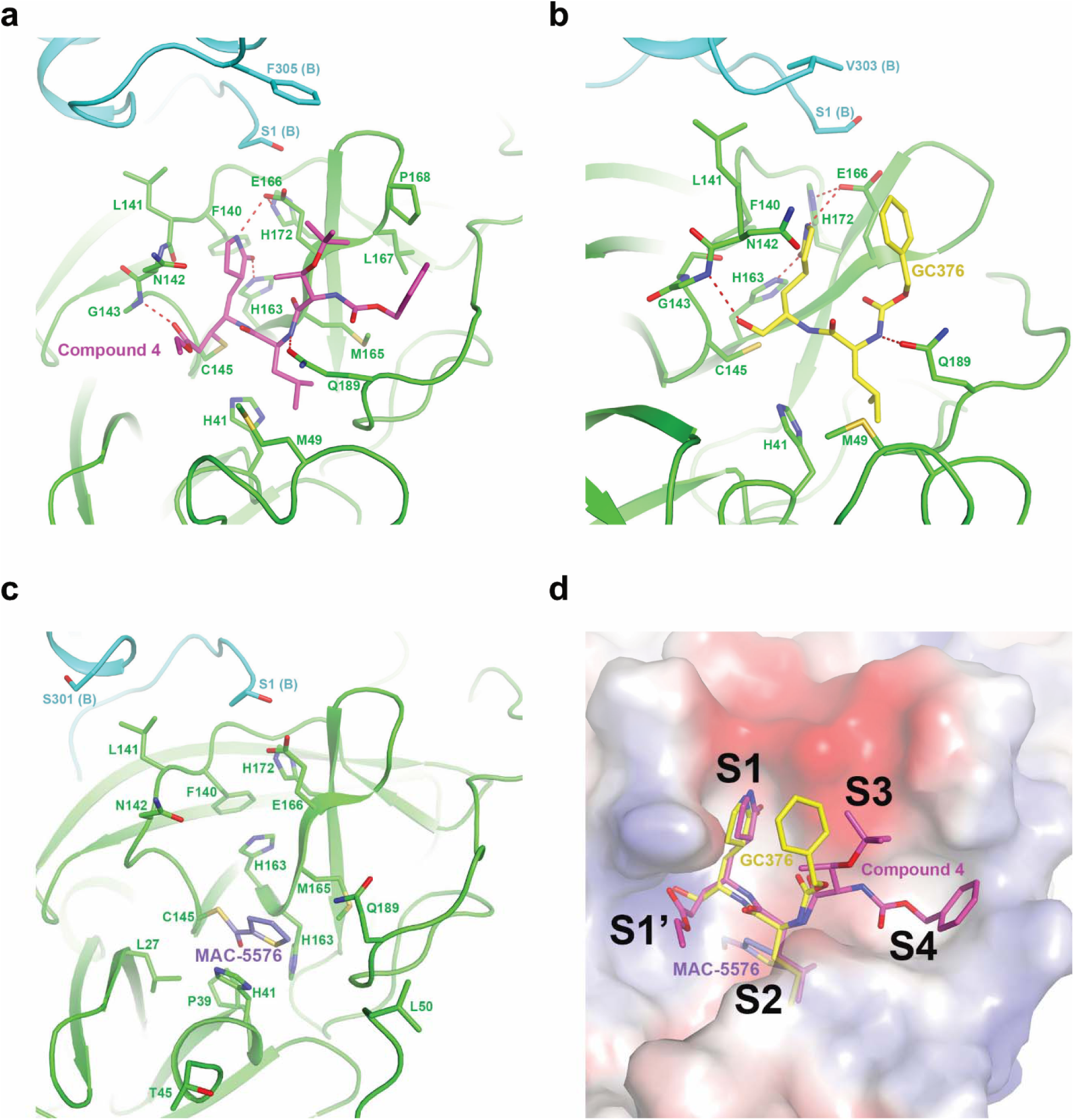
Crystal structures of inhibitors in complex with the SARS-CoV-2 3CL protease. **a-c**, Structure of compound 4 (**a**), GC376 (**b**), or MAC-5576 (**c**) bound to SARS-CoV-2 3CL. Protomer A is denoted in green and protomer B is denoted in cyan. **d**, Overlay of all three compounds in the substrate binding pocket of the 3CL protease.

The X-ray crystal structures revealed that all three of the compounds bind covalently to the catalytically active Cys145 residue within the substrate-binding pocket of the protease. We observed distinct mechanisms by which these compounds acted on this residue. Compound 4 functioned in a similar binding mode as other reported compounds, covalently modifying Cys145 through Michael addition (**Fig. 2a**)^5^. For GC376, the bisulfite adduct was converted to an aldehyde as previously reported, allowing it to then react with Cys145 through nucleophilic addition and hemithioacetal formation (**Fig. 2b**)^8^. MAC-5576 also covalently modified Cys145 by nucleophilic linkage, as expected (**Fig. 2c**).

As we solved the structures for multiple compounds, we hypothesized that general principles for the design of SARS-CoV-2 3CL protease inhibitors could be identified. We first overlaid all four crystal structures of the 3CL with or without inhibitors (**Extended Data Fig. 5**). We observed local conformational changes, with Thr45 to Pro52 distinct from the *apo* 3CL in all three inhibitor-bound structures, whereas Arg188 to Gln192 differed only in the compound 4 and GC376-bound, but not MAC-5576-bound structures. We then overlaid each of the inhibitors in the substrate binding pocket of the 3CL protease to find commonalities in their interactions (**Fig. 2d**). Most notably, we found that all of these compounds occupied the S2 site, with compound 4 and GC376 further anchored in the S1 and S3 sites. The backbone NH of Gly143 points toward the ligand binding pocket, forming hydrogen bonds with the carbonyl oxygen of the ethyl ester of compound 4, and the hemithioacetal of GC376 after the Cys145 addition to the original aldehyde, even though the former hydrogen bond is stronger than the latter. In both structures, the γ-lactam groups occupy the S1 site, and are strongly anchored by two hydrogen bonds with the side chains of His163 and Glu166. The isobutyl groups are favorably embedded in the hydrophobic S2 site, surrounded by the alkyl portion of the side chains of His41, Met49, His164, Met165, Asp187, and Gln189. Extending into the S3 pocket, the amide bonds of compound 4 and GC376 are stabilized by hydrogen bond interactions with the side chain of Gln189. Similar interactions are also observed in reports of related compounds, suggesting that overall, the binding modes of this class of substrate mimetic inhibitors share remarkable similarities^5,6,10^. Specifically, they all have a γ-lactam occupying the S1 pocket, preserving the dual hydrogen bonds with His163 and Glu166. Furthermore, they commonly contain a hydrophobic moiety occupying the S2 site. As shown in a structural overlay of compound 4 and GC376 with these related compounds (**Extended Data Fig. 6**), the segment of the inhibitors from S1 to S2 align closely on top of each other. Variations of binding start to emerge in the S3 and S4 region, which exhibits high degrees of freedom in terms of structural diversity as well as conformational flexibility. In our experiments, the S3 and S4 sites displayed weaker electron density, indicating flexibility in the inhibitor and/or the protease in these regions (**Extended Data Fig. 4**). These observations suggest that development of 3CL protease inhibitors may benefit from first establishing robust interactions within the S1, S2, and/or S1’ sites, before extending into the S3 and S4 sites. Possibly, compounds such as compound 4 and GC376 are not optimized for binding into the S3 and S4 sites, and there are ample opportunities to improve the inhibitory potencies against the 3CL by designing compounds that exploit the accessible contact points to strengthen the ligand-protein interactions.

On the other hand, the binding of MAC-5576, as a non-peptidic small molecule, displays unique features that differ from that of compound 4 or GC376. We observed that the thiophene group forms π-π stacking with the His41 side chain imidazole, which undergoes a conformational rotation around its beta-carbon to align parallel to the thiophene, as compared to the other peptide-bound structures. Additionally, the side chain of Gln189 also shows notable conformational variation compared to those in the compound 4 and GC376 crystal structures, possibly in response to the specific hydrogen bond interactions induced by the respective ligands. Notably, the rotation of His41 has been reported previously in the crystal structure of a benzotriazole ester inhibitor (XP-59) in complex with the SARS-CoV-1 3CL protease (PDB:2V6N)^11^. An overlaid model of the crystal structures of MAC-5576 bound to SARS-CoV-2 3CL and XP-59 bound to SARS-CoV-1 3CL shows that both compounds have similar binding modes when covalently bound to Cys145, in which the thiophene of MAC-5576 and the phenyl ring of XP-59 almost overlap with each other, both engaging the His41 side chain via π-π stacking interactions (**Extended Data Fig. 7**).

In summation, we have identified compound 4, GC376, and MAC-5576 as inhibitors of the SARS-CoV-2 3CL protease. Crystal structures of the compounds complexed to the protease suggested their mechanisms of action, as well as portended guidelines for the development of SARS-CoV-2 3CL protease inhibitors, which may aid in the future development of novel inhibitors to combat this virus.

## Methods

### Compounds

Compound 4 was synthesized using the synthesis route previously described, with the exception of using a sodium borohydride-cobaltous chloride reduction of the nitrile in the construction of the lactam, thus avoiding the high pressure hydrogenation in the original route^7, 12^. GC376 was purchased from Aobious and MAC-5576 was purchased from Maybridge.

### Expression and purification of SARS-CoV-2 3CL protease

The SARS-CoV-2 3CL protease gene was codon optimized for bacterial expression and synthesized (Twist Bioscience), then cloned into a bacterial expression vector (pGEX-5X-3, GE) that expresses the protease as a fusion construct with a N-terminal GST tag, followed by a Factor Xa cleavage site. After confirmation by Sanger sequencing, the construct was transformed into BL21 (DE3) cells. These *E. coli* were inoculated and grown overnight as starter cultures, then used to inoculate larger cultures at a 1:100 dilution, which were then grown at 37 °C, 220 RPM until the OD reached 0.6-0.7. Expression of the protease was induced with the addition of 0.5 mM IPTG, and then the cultures were incubated at 16 °C, 180 RPM for 10 h. Cells were pelleted at 4500 RPM for 15 min at 4 °C, resuspended in lysis buffer (20 mM Tris-HCl, pH 8.0, 300 mM NaCl), homogenized by sonication, then clarified by centrifuging at 25000 x *g* for 1 h at 4 °C. The supernatant was mixed with Glutathione Sepharose resin (Sigma) and placed on a rotator for 2 h at 4 °C. The resin was then repeatedly washed by centrifugation at 3500 RPM for 15 min at 4 °C, discarding of the supernatant, and then resuspension of the resin in fresh lysis buffer. After ten washes, the resin was resuspended in lysis buffer, and Factor Xa was added and incubated for 18 h at 4 °C on a rotator. The resin was centrifuged at 3500 RPM for 15 min at 4 °C, and then the supernatant was collected and concentrated using a 10 kDa concentrator (Amicon) before being loaded onto a Superdex 10/300 GL column for further purification by size exclusion chromatography. The appropriate fractions were collected and pooled with a 10 kDa concentrator, and then the final product was assessed for quality by SDS-PAGE and measurement of biochemical activity.

### Measurement of SARS-CoV-2 3CL protease biochemical activity

The *in vitro* biochemical activity of the SARS-CoV-2 3CL protease was measured as previously described^5^. The fluorogenic peptide MCA-AVLQSGFR-Lys(DNP)-Lys-NH2, corresponding to the nsp4/nsp5 cleavage site in the virus, was synthesized (GL Biochem), then resuspended in DMSO to use as the substrate. Different concentrations of this substrate, ranging from 5 µM to 100 µM, were prepared in the assay buffer (50 mM Tris-HCl, pH 7.5, 1 mM EDTA) in a 96 well-plate. The protease was then added to each well at a concentration of 0.2 µM, and then fluorescence was continuously measured on a plate reader for 3 min. The catalytic efficiency of the protease was then calculated by generating a double-reciprocal plot.

### Measurement of SARS-CoV-2 3CL protease inhibition

Inhibition of the biochemical activity of the SARS-CoV-2 3CL protease was quantified as previously described with modifications^5^. Serial dilutions of the test compound were prepared in the assay buffer, and then incubated with 0.2 µM of the protease for 10 min at 37 °C. The substrate was then added at 20 µM per well, and then fluorescence was continuously measured on a plate reader for 3 min. Inhibition was then calculated by comparison to control wells with no inhibitor added. IC_50_ values were determined by fitting an asymmetric sigmoidal curve to the data (GraphPad Prism).

### Measurement of SARS-CoV-2 viral inhibition

Stocks of SARS-CoV-2 strain 2019-nCoV/USA_WA1/2020 was propagated and titered in Vero-E6 cells. One day prior to the experiment, Vero-E6 cells were seeded at 30,000 cells/well in 96 well-plates. Serial dilutions of the test compound were prepared in cell media (EMEM + 10% FCS + penicillin/streptomycin), overlaid onto cells, and then virus was added to each well at an MOI of 0.2. Cells were incubated at 37 °C under 5% CO_2_ for 72 h and then viral cytopathic effect was scored in a blinded manner. Inhibition was calculated by comparison to control wells with no inhibitor added. EC_50_ values were determined by fitting an asymmetric sigmoidal curve to the data (GraphPad Prism). Cells were confirmed as mycoplasma negative prior to use. All experiments were conducted in a biosafety level 3 (BSL-3) lab.

### Measurement of cellular cytotoxicity

Vero-E6 cells were incubated with the compound of interest for 48 h at 37 °C under 5% CO_2_ and then cellular cytotoxicity was determined with the XTT Cell Proliferation Assay Kit (ATCC) according to the manufacturer’s instructions.

### Crystallization, data collection, and structure determination

To generate the complex of SARS-CoV-2 3CL protease bound to compound 4, 50 µM of the 3CL protease was incubated with 500 µM of compound 4 in a buffer comprised of 50 mM Tris-HCl (pH 7.5), 1 mM EDTA, and 5% (v/v) glycerol for 1 h at 4 °C. This complex was then concentrated to 8.5 mg/mL using a 10 kDa concentrator, and initially subjected to extensive robotic screening at the High-Throughput Crystallization Screening Center^13^ of the Hauptman-Woodward Medical Research Institute (HWI) (https://hwi.buffalo.edu/high-throughput-crystallization-center/). The most promising crystal hits were then reproduced using the microbatch-under-oil method at 4 °C. Block-like crystals of 3CL in complex with compound 4 appeared after a few days in the crystallization condition comprised of 0.1 M potassium nitrate, 0.1 M sodium acetate (pH 5), and 20% (w/v) PEG 1000 with protein to crystallization reagent at a 2:1 ratio. The crystals were subsequently transferred into the same crystallization reagent supplemented with 15% (v/v) glycerol and flash-frozen in liquid nitrogen.

To obtain crystals of 3CL in complex with GC376, crystals of *apo* 3CL were initially grown by using seeding method in a crystallization reagent comprised of 0.1 M sodium phosphate-monobasic, 0.1 MES (pH 6), and 20% (w/v) PEG 4000. The *apo* crystals were subsequently soaked with 15 Mm GC376, followed by flash-freezing of the crystals in the same reagent supplemented with 15% ethylene glycol.

To generate the complex of SARS-CoV-2 3CL protease bound to MAC-5576, 50 µM of the 3CL protease was incubated with 500 µM of compound 4 in a buffer comprised of 50 mM Tris-HCl (pH 7.5), 1 mM EDTA, and 5% (v/v) glycerol for 1 h at 4 °C. The complex was concentrated to 10 mg/mL using a 10 kDa concentrator, and then crystallized in the same conditions as those used for crystallization of *apo* 3CL.

A native dataset was collected on each crystal of 3CL, alone (*apo*), and in complex with compound 4 and GC376 at the NE-CAT24-ID-C beam line of Advanced Photon Source (APS) at Argonne National Laboratory, and the NE-CAT 24-ID-E beam line of APS was used for data collection on crystals of 3CL-MAC-5576. Crystals of *apo* 3CL and in complex with compound 4, GC376, and MAC-5576 diffracted the X-ray beam to resolution 1.85 Å, 1.80 Å, 1.83 Å, 1.73 Å, respectively. The images were processed and scaled in space group C2 using XDS^14^. The structure of 3CL-compound 4 was determined by molecular replacement method using program MOLREP^15^ and the crystal structure of 3CL in complex with inhibitor N3 (PDB id: 6LU7)^5^ was used as a search model. The geometry of each crystal structure was subsequently fixed and the corresponding inhibitor was modeled in by XtalView^16^ and Coot^17^, and refined using PHENIX^18^. The mapping of electrostatic potential surfaces was generated in PyMOL with the APBS plug-in^19^. There is one protomer of 3CL complex in the asymmetric unit of each crystal. The crystallographic statistics are shown in **Supplementary Table 1**.

### Reporting Summary

Further information on research design is available in the Nature Research Reporting Summary linked to this article.

## Data availability

Structural data for the *apo* SARS-CoV-2 3CL protease and 3CL in complex with compound 4, GC376, and MAC-5576 will be deposited in the Protein Data Bank (PDB) and made publicly available upon publication. Source data for Fig. 1, Extended Data Fig. 1b, Extended Data Fig. 2, and the unprocessed gel for Extended Data Fig. 1a are available with the paper online.

## Acknowledgements

This work was supported by a grant from the Jack Ma Foundation to D.D.H. and A.C. and by grants from Columbia Technology Ventures and the Columbia Translational Therapeutics (TRx) program to B.R.S. A.C. is also supported by a Career Awards for Medical Scientists from the Burroughs Wellcome Fund. S.I. is supported by NIH grant T32AI106711. We thank the staff of the High-Throughput Crystallization Screening Center of the Hauptman-Woodward Medical Research Institute for screening of crystallization conditions and the staff of the Advanced Photon Source at Argonne National Laboratory for assistance with data collection.

## Author contributions

D.D.H. conceived the project. S.I., B.R.S., A.C., and D.D.H. planned and designed the experiments. S.I. and S.J.H. cloned, expressed, and purified proteins. A.Z. synthesized compound 4. S.I. conducted the protease inhibition assay. M.N. and Y.H. conducted the antiviral assay and cytotoxicity measurements. F.F. and H.L. crystallized the proteins. F.F. collected diffraction data and solved the crystal structures. F.F., F.-Y.L., and L.X. analyzed the structural data. S.I., F.F., F.-Y.L., L.X., A.C., and D.D.H. wrote the manuscript with input from all authors.

---

**Extended Data Fig. 1.**
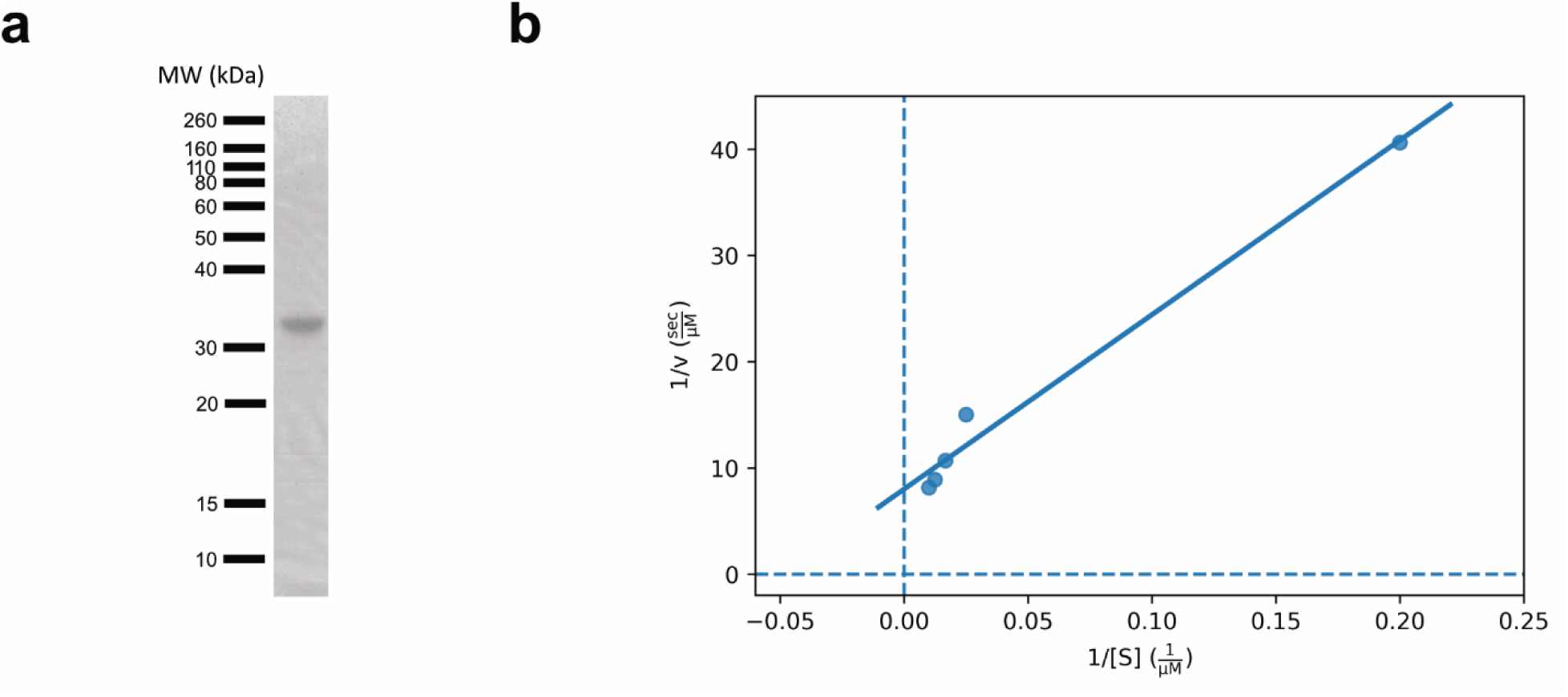
Production of native SARS-CoV-2 3CL protease in E. coli. **a**, The purified protease was ran on SDS-PAGE to confirm size and purity. **b**, Confirmation of enzymatic activity of SARS-CoV-2 3CL protease by quantification of cleavage of a fluorogenic peptide substrate.

**Extended Data Fig. 2.**
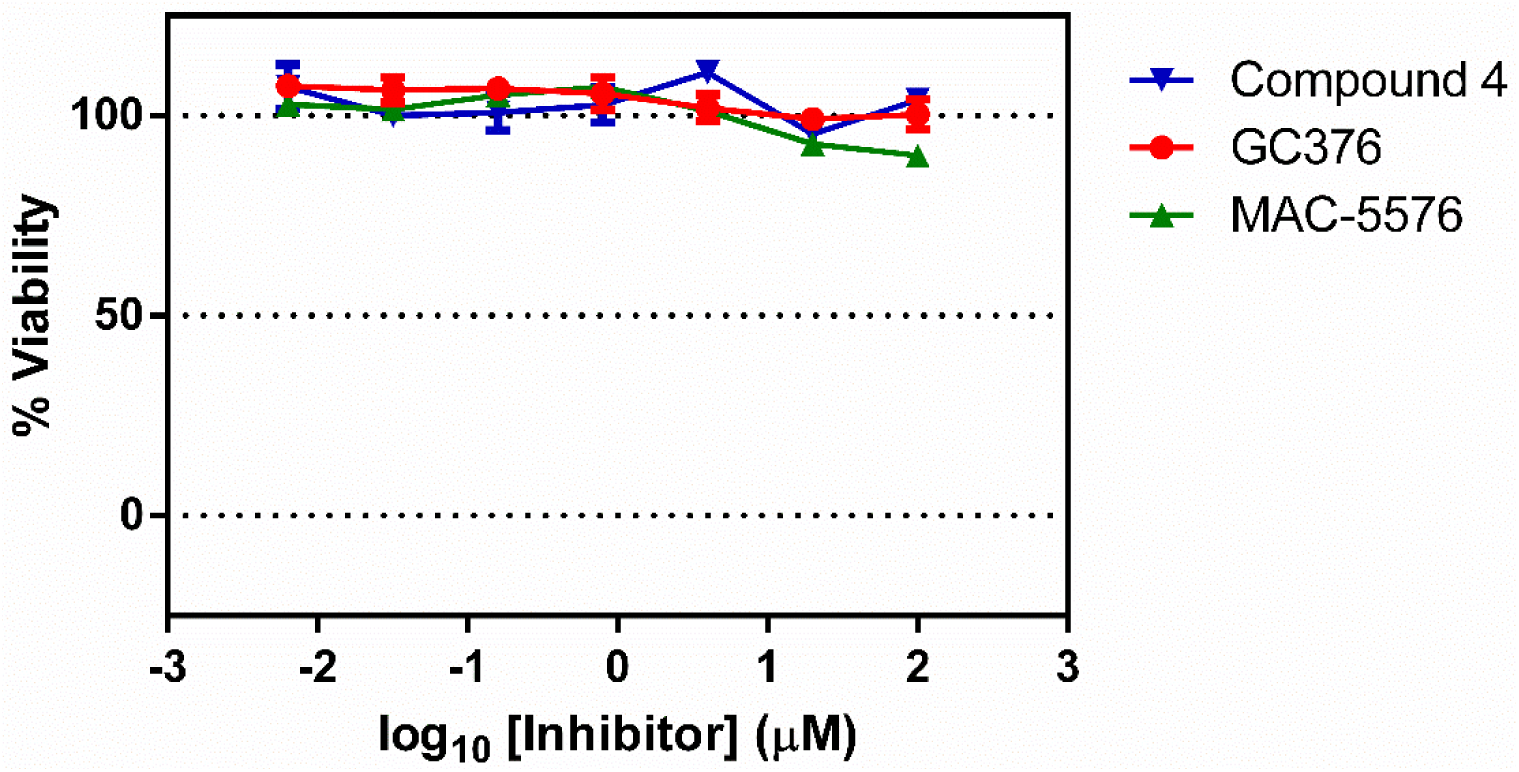
Cytotoxicity of compounds in Vero-E6 cells. Vero-E6 cells were incubated with serial dilutions of each compound for 48 h and then tested for cytotoxicity. Data are shown as mean ± s.e.m. for three experimental replicates.

**Extended Data Fig. 3.**
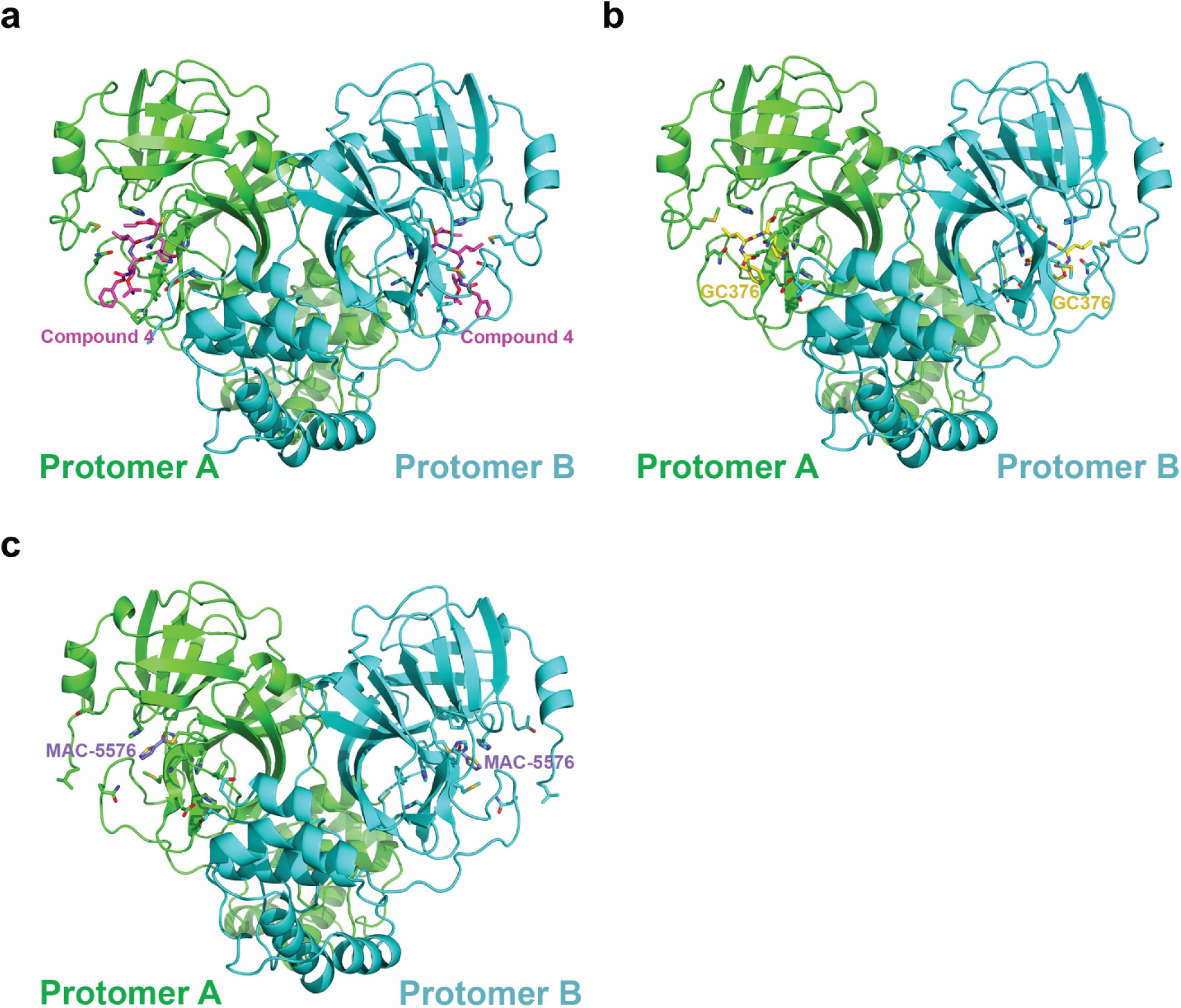
Overall crystal structures of inhibitors in complex with the SARS-CoV-2 3CL protease dimer. **a-c**, Structures of compound 4 (**a**), GC376 (**b**), or MAC-5576 (**c**) bound to the 3CL dimer.

**Extended Data Fig. 4.**
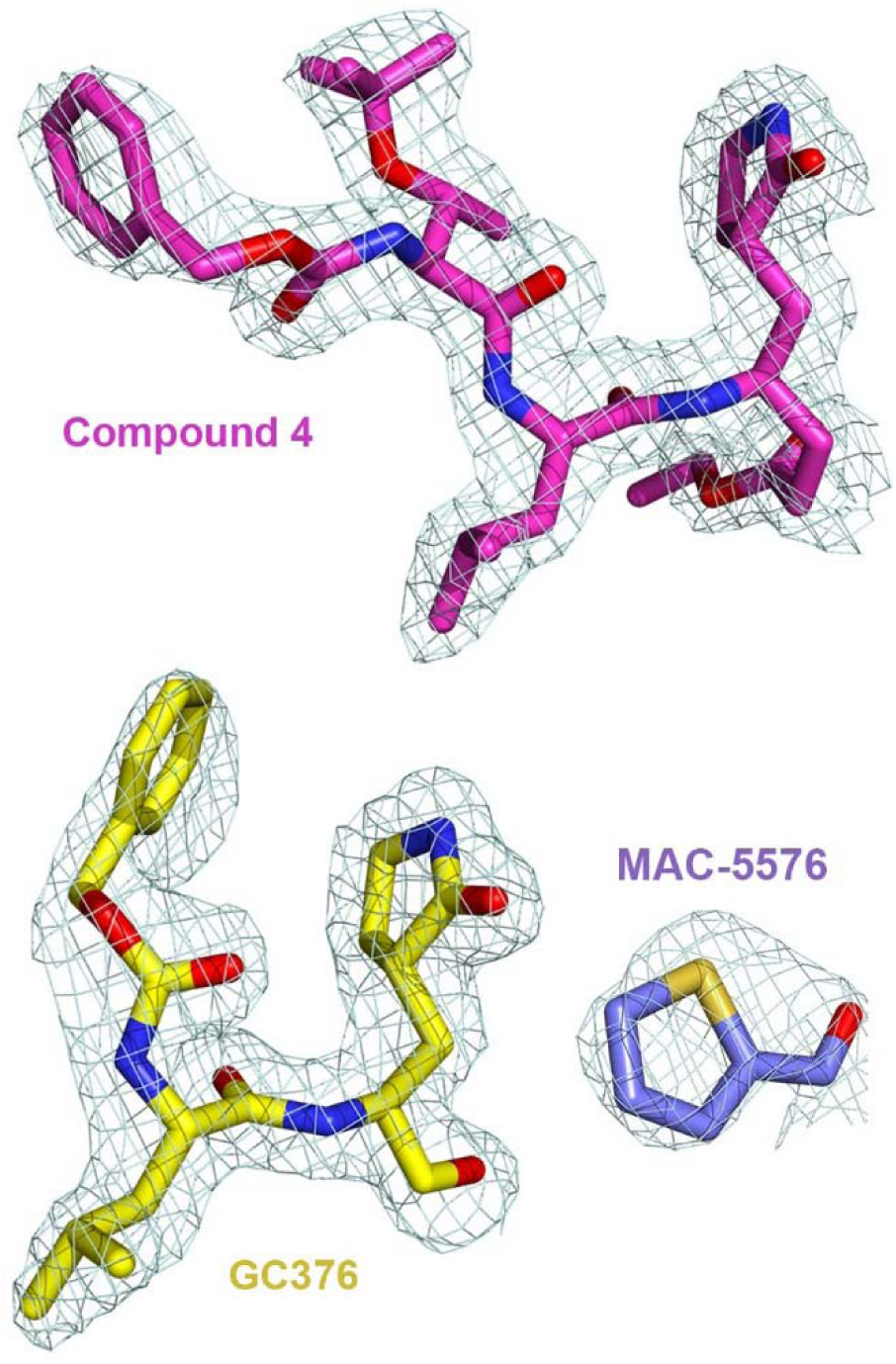
2Fo–Fc electron density map at 1σ for the inhibitors in the crystal structures. Electron density mesh (light cyan) for compound 4, GC376, and MAC-5576 in complex with 3CL.

**Extended Data Fig. 5.**
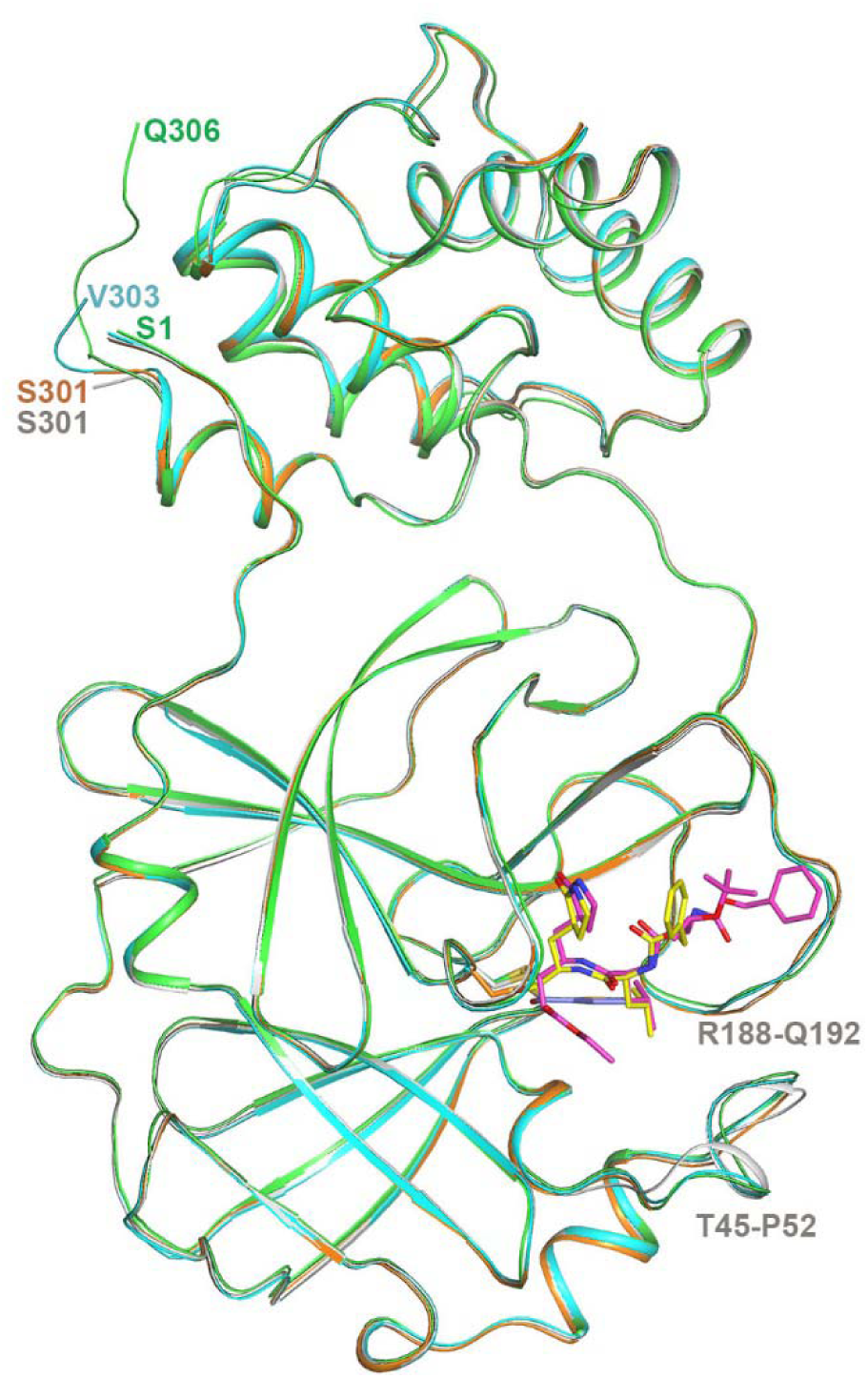
Overlay of the four crystal structures of 3CL. Crystal structure of 3CL in *apo* form (gray), in complex with, compound 4 (green for 3CL and magenta for compound 4), GC376 (cyan for 3CL and yellow for GC376), and MAC-5576 (orange for 3CL and purple for MAC-5576 fragment). One protomer for each structure is shown, with the inhibitors shown with stick models. The terminal residue of each structure, as well as two stretches of residues near the binding site that exhibit local conformational change between the *apo* and inhibitor-bound structures are labeled.

**Extended Data Fig. 6.**
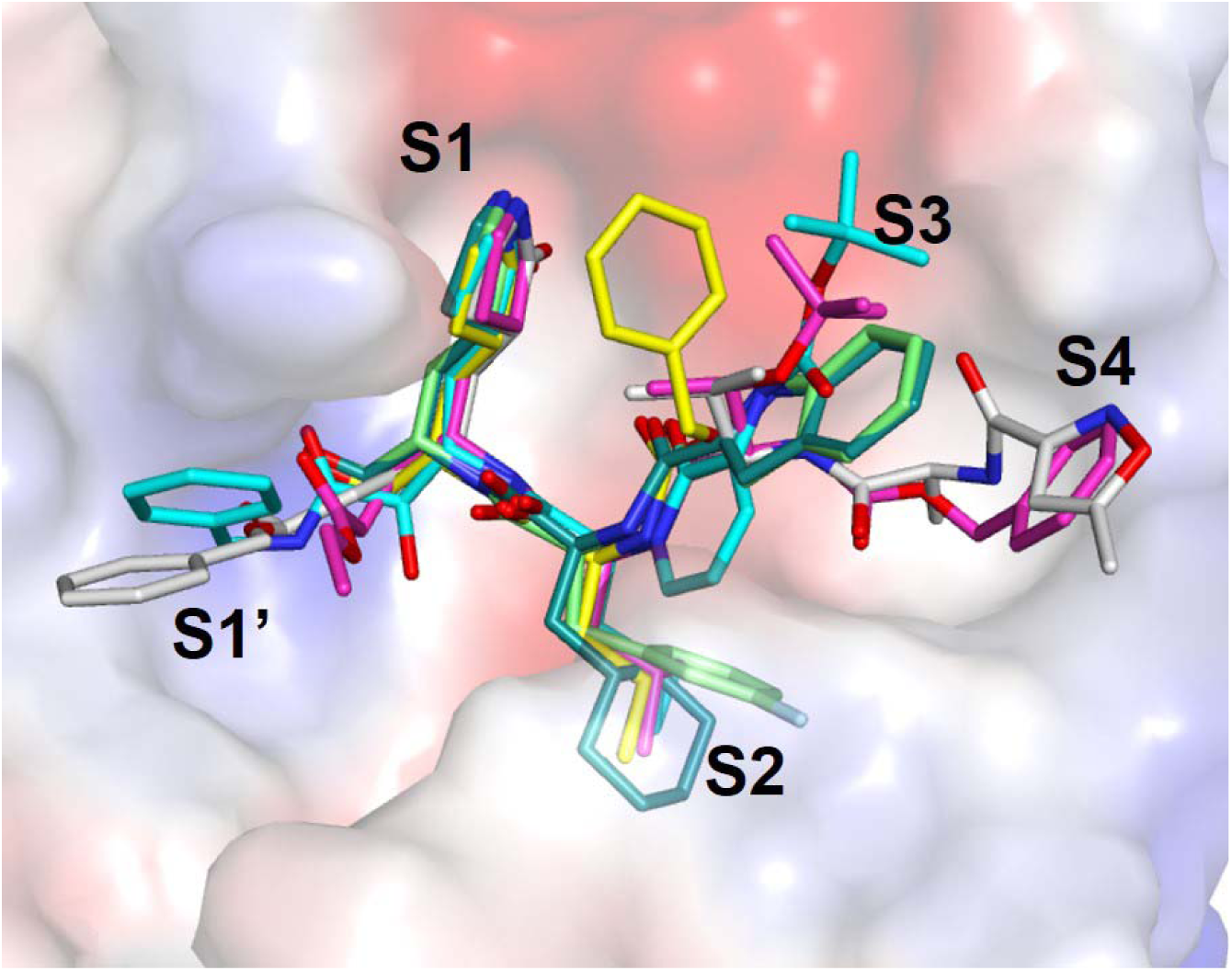
Comparison of the binding modes of compound 4 and GC376 with other peptide-like inhibitors. Compound 4 (magenta) and GC376 (yellow) were overlaid with previously identified compounds, compound 13b (cyan, PDB: 6Y2F), compound 11a (dark green, PDB: 6LZE), compound 11b (light green, PDB: 6M0K), and N3 (white; PDB: 7BQY).

**Extended Data Fig. 7.**
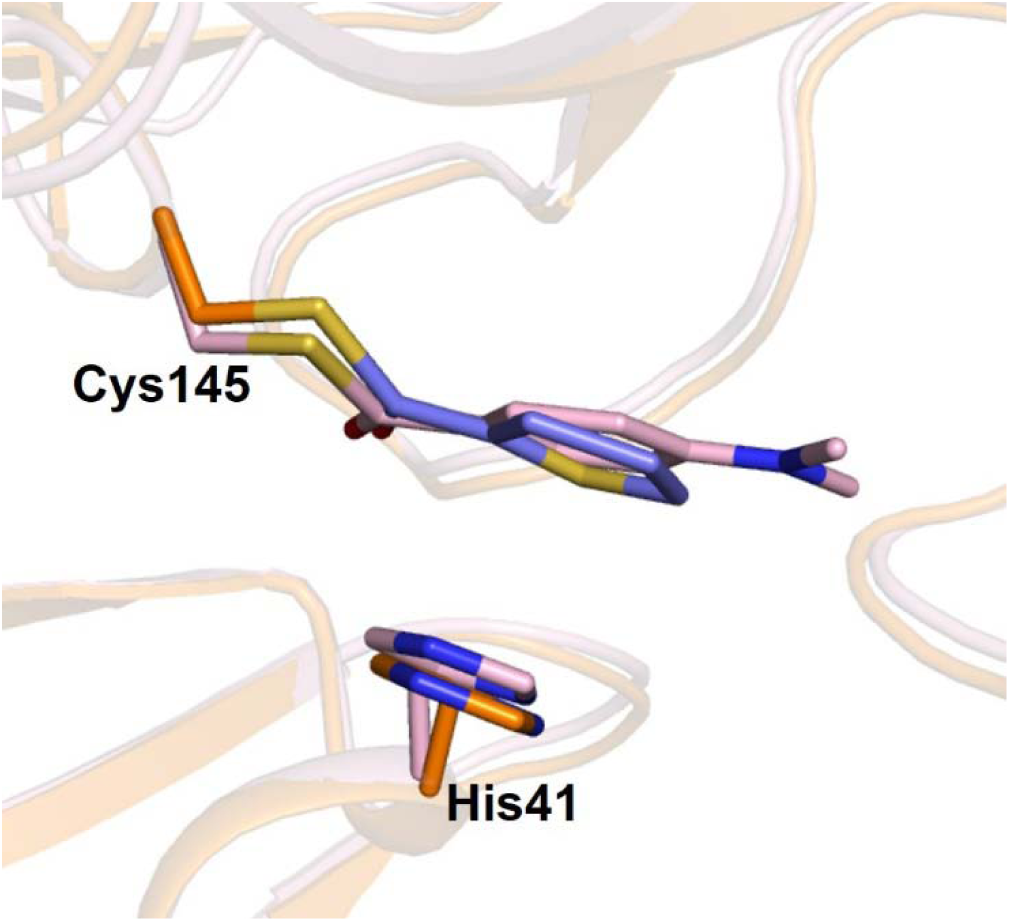
Comparison of the binding modes of MAC-5576 with XP-59. MAC-5576 (purple) bound to the SARS-CoV-2 3CL protease (orange) was overlaid with XP-59 bound to the SARS-CoV-1 3CL protease (pink, PDB: 2V6N).

**Supplemental Table 1.** Data collection and refinement statistics of *apo* SARS-CoV-2 3CL protease and 3CL protease bound to compound 4, GC376, and MAC-5576.

|  | *apo* 3CL | 3CL with<br>Compound 4 | 3CL with<br>GC376 | 3CL with<br>MAC-5576 |
| --- | --- | --- | --- | --- |
| **Data collection** |  |  |  |  |
| Space group | *C*2 | *C*2 | *C*2 | *C*2 |
| Cell dimensions |  |  |  |  |
| *a*, *b*, *c* (Å) | 98.7, 82.0, 51.8 | 97.2, 81.9, 54.2 | 98.8, 80.2, 52.0 | 98.3, 82.5, 51.8 |
| $\alpha$, $\beta$, $\gamma$ (°) | 90, 114.9, 90 | 90, 117.1, 90 | 90, 114.3, 90 | 90, 114.83, 90 |
| Resolution (Å) | 60.4-1.85<br>(1.88-1.85)* | 59.5-1.80<br>(1.82-1.80)* | 59.9-1.83<br>(1.86-1.83)* | 60.58-1.73<br>(1.76-1.73)* |
| *R*<sub>merge</sub> (%) | 7.2 (65.7) | 18.7 (57.6) | 14.7 (60.5) | 4.1 (68.8) |
| *I*/σ*I* | 13.4 (2.1) | 10.2 (2.4) | 12.7 (2.2) | 23.5 (2.3) |
| Completeness (%) | 98.9 (97.4) | 98.8 (99.0) | 98.5 (96.0) | 99.2 (88.3) |
| Redundancy | 6.8 (6.5) | 6.7 (6.5) | 6.9 (6.1) | 6.8 (6.6) |
| CC1/2 | 99.8 (95.1) | 0.99 (0.91) | 0.99 (0.93) | 100 (0.90) |
| **Refinement** |  |  |  |  |
| Resolution (Å) | 47.0-1.85<br>(1.88-1.85)* | 48.2-1.80<br>(1.82-1.80)* | 47.38-1.83<br>(1.86-1.83) | 47.0-1.73<br>(1.75-1.73) |
| No. reflections | 31,710 (3,219) | 34,294 (3,497) | 31,972 (3,134) | 38,879 (3,899) |
| *R*<sub>work</sub> / *R*<sub>free</sub> (%) | 16.8 (25.2)/19.8<br>(29.3) | 18.7 (25.7)/22.3<br>(30.7) | 17.4 (25.0)/20.6<br>(28.7) | 16.6 (26.8)/19.0<br>(27.6) |
| No. atoms |  |  |  |  |
| Protein | 2,329 | 2,370 | 2,340 | 2,298 |
| Ligand/ion | 5 | 44 | 28 | 16 |
| Water | 128 | 293 | 163 | 245 |
| B-factors |  |  |  |  |
| Protein | 50.0 | 35.2 | 41.4 | 40.5 |
| Ligand/ion | 52.6 | 36.6 | 42.4 | 43.7 |
| Water | 53.5 | 46.7 | 48.2 | 50.7 |
| R.m.s deviations |  |  |  |  |
| Bond lengths (Å) | 0.006 | 0.007 | 0.006 | 0.006 |
| Bond angles (°) | 0.8 | 0.8 | 0.8 | 0.8 |

\*Highest resolution shell is shown in parenthesis.

